# Effects of Orientation Change during Environmental Learning on Age-Related Difference in Spatial Memory

**DOI:** 10.1101/412965

**Authors:** Naohide Yamamoto, Michael J. Fox, Ellen Boys, Jodi Ord

**Affiliations:** School of Psychology and Counselling, Queensland University of Technology, Kelvin Grove, QLD 4059, Australia; Institute of Health and Biomedical Innovation, Queensland University of Technology, Kelvin Grove, QLD 4059, Australia; Department of Psychology, Cleveland State University, Cleveland, OH 44115, USA

**Keywords:** aging, medial temporal lobe, navigation, map, route, survey

## Abstract

It has been suggested that older adults suffer a greater degree of decline in environmental learning when navigating in an environment than when reading a map of the environment. However, the two types of spatial learning differ not only in perspectives (i.e., navigation is done with a ground-level perspective; a map is read from an aerial perspective) but also in orientations (i.e., orientations vary during navigation; spatial information is drawn from a single orientation in a map), making it unclear which factor critically affects older adults’ spatial learning. The present study addressed this issue by having younger and older participants learn the layout of a large-scale environment through an aerial movie that contained changes in orientations from which the environment was depicted. Results showed that older participants’ memories for the environmental layout were as distorted as those created through a ground-level movie (which involved the same orientation changes), whereas they formed more accurate memories through another aerial movie in which an orientation was fixed. By contrast, younger participants learned the environment equally well from the three movies. Taken together, these findings suggest that there is age-related alteration specifically in the ability to process multiple orientations of an environment while encoding its layout in memory. It is inferred that this alteration stems from functional deterioration of the medial temporal lobe, and possibly that of posterior cingulate areas as well (e.g., the retrosplenial cortex), in late adulthood.

## 1. Introduction

The ability to learn an environmental layout is susceptible to adverse effects of aging. Not only those who have dementia but also some of the healthy elderly experience difficulty in spatial learning and navigation in a large-scale environment [1–3]. This age-related change may be in part due to deficits of general cognitive functions such as attention and working memory [4]. However, not all spatial and navigational skills show the same pattern of decline in senescence [5–7], suggesting that global factors do not fully characterize the way in which younger and older adults learn spatial layouts differently.

To gain insights into how different types of spatial learning are differentially affected by aging, Yamamoto and DeGirolamo [7] examined healthy older adults’ spatial memories after they viewed movies of large-scale environments taken from two perspectives: a ground-level view of an observer navigating in an environment (Figure 1A) and an aerial view of another observer looking straight down a section of the environment (Figure 1B). The movies simulated spatial learning by first-person navigation and map reading, respectively [8]. It was hypothesized that the former would be more susceptible to the effects of aging than the latter because (a) the medial temporal lobe (MTL) including the hippocampus tends to be more heavily involved in encoding environmental layouts from the ground-level perspective than the aerial perspective [9–11] and (b) age-related atrophy is particularly notable in the MTL and the reduction of the MTL volume seems to be a factor in the deterioration of spatial learning and memory in old age [12–17]. Yamamoto and DeGirolamo indeed found that, relative to the baseline level of performance shown by younger adults, older adults’ memories for landmark locations in the environments were more distorted when they were learned through the ground-level movie than the aerial movie. That is, the older adults exhibited a greater degree of decline of spatial learning via first-person navigation than via map reading, lending clear support to the hypothesis.

**Figure 1.**
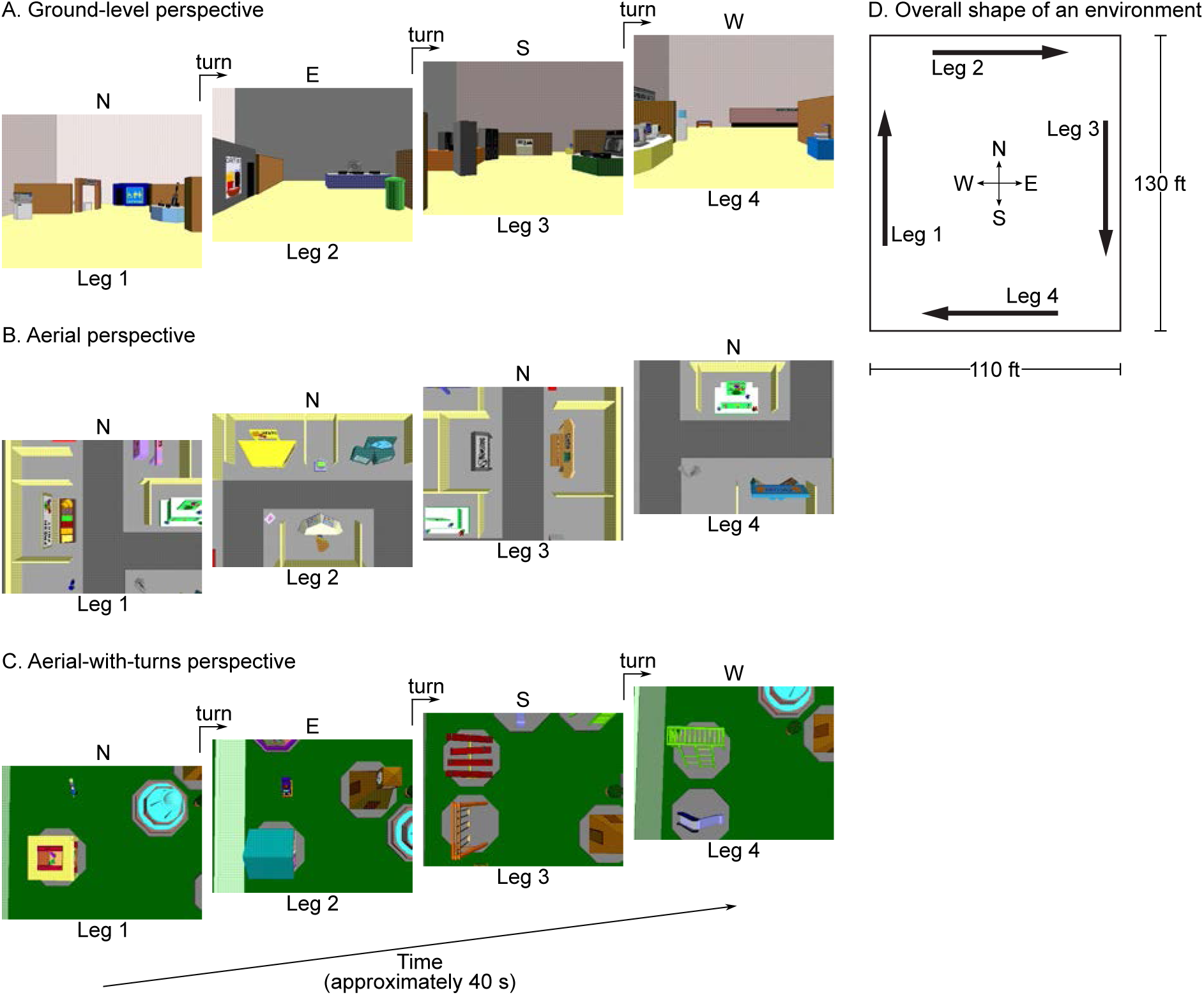
Three perspectives from which virtual environments were presented in the experiment. Letters above each snapshot of an environment indicate heading directions (N = north; E = east; S = south; W = west). In the ground-level and aerial-with-turns perspectives, there were 90° turns to the right at each corner, leading to changes in participants’ orientations within the environment. In the aerial perspective, participants maintained the north-up orientation all the time. For example, in leg 2, participants’ view moved from left to right along the leg while holding the northerly heading constant. These virtual environments were originally created and used by Shelton and colleagues [8, 10, 11].

This finding, however, did not entirely clarify why spatial learning through first-person navigation was particularly challenging to older adults [7]. This unclarity stemmed from the fact that the ground-level and aerial movies differed not only in their perspectives but also in the variation of participants’ orientations (or lack thereof) within environments. In the ground-level movie, participants experienced four different orientations of an environment as they turned each corner (Figure 1A). On the other hand, in the aerial movie, participants always viewed the environments in the north-up orientation (Figure 1B). Importantly, this difference was not a mere experimental artifact; rather, it reflected the different ways observers learn environmental layout by actually navigating in the environment (during which they are exposed to varying orientations) and by reading a map (which typically presents spatial information using a fixed orientation). Nevertheless, the design of the movies made it unclear whether it was the ground-level perspective itself or the exposure to multiple orientations that caused the greater extent of age-related decline in spatial learning in the ground-level condition.

The present study was designed to address this issue by contrasting two hypotheses. One hypothesis was that learning an environmental layout from a ground-level perspective could be more challenging than learning the same layout from an aerial perspective, especially for older adults (*perspective hypothesis*). When an environment is viewed from the ground-level perspective, some information about its layout is not immediately available in the first-person view. Instead, it must be reconstructed through inferential processes. For example, distances between landmarks appear differently depending on their orientations: Intervals of the same length are perceived to be shorter when they extend in depth than when they run parallel to observers’ frontal plane [18–20]. As a result, learning the layout of landmarks from the ground-level perspective entails additional mental computation for correcting for this perceptual bias, which is not necessary when all intervals between landmarks are placed on the same plane in an aerial view. Older adults can be proficient at counteracting these biases, presumably due to accumulated experience in their lifetime, when they have sufficient time to observe a static environment [21–23]. However, given that the speed with which cognitive operations are carried out is reduced overall in late adulthood [24, 25], older adults might be disadvantaged when they need to swiftly extract spatial information from dynamic images of the environment during first-person navigation.

The other hypothesis was that viewing an environment from multiple orientations would increase the difficulty of learning its layout (*orientation hypothesis*). When the to-be-learned environment is large-scale, not all landmarks are visible from a single location. As a result, unless a navigator encodes all landmark locations using an allocentric (i.e., environment-centered) frame of reference, landmarks viewed from different orientations are initially represented in different egocentric frames of reference. When forming a memory for the entire layout of the environment, some landmark pairs need to be mentally rotated so that all landmarks are placed within a common frame of reference. It is possible that a greater degree of distortion is introduced in older adults’ spatial memory through this process because they perform mental rotation with reduced speed and accuracy compared to younger adults [26–29]. On the other hand, when the environment is learned from an aerial perspective in a fixed orientation, spatial relations between landmarks can be encoded in the same frame of reference from the beginning. Landmarks that are not viewed together still need to be integrated to represent the whole layout in memory, but this integration does not require mental rotation, making it less likely that substantial age-related difference arises from this operation.

To test the perspective and orientation hypotheses, the present study introduced a new condition in the procedure of the Yamamoto and DeGirolamo [7] study: In addition to the ground-level and aerial movies, participants viewed a movie in which an environment was shown from an aerial perspective but with varying orientations as in the ground-level movie (Figure 1C). In the new movie, the entire environment was rotated by 90° at each corner so that participants experienced the four legs of the environment in different orientations. This aerial-with-turns movie was effective for distinguishing between the hypotheses because different patterns of spatial learning performance were predicted under the two hypotheses. The perspective hypothesis posits that the primary source of older adults’ disadvantage in spatial learning would not be variation of orientations but the ground-level perspective. Because the aerial-with-turns and original aerial movies share the aerial perspective, it follows that these aerial movies should yield equivalent spatial memories, which would be more accurate than spatial memories originating from the ground-level movie. On the other hand, the orientation hypothesis postulates that exposure to multiple orientations would be the critical factor that makes spatial learning via the ground-level movie particularly difficult for older adults. According to this hypothesis, the aerial-with-turns movie should function in the same manner as the ground-level movie because these movies involve the identical changes in orientation. Spatial memories that result from the aerial-with-turns and ground-level movies would be less accurate than those from the original aerial movie.

Although both of the hypotheses are theoretically plausible, empirical evidence lends stronger support to the orientation hypothesis. Using functional magnetic resonance imaging (fMRI), Shelton and Pippitt [11] investigated neuronal activation while participants learned large-scale environments through the three types of movies. There were two important findings in this study. First, spatial learning through the ground-level movie elicited greater bilateral activation in the MTL than spatial learning through the aerial movie in a fixed orientation, replicating previous results [10]. Second, more critically for the present study, the MTL was bilaterally activated to the same extent by the ground-level movie and the aerial-with-turns movie, which was greater than the level of MTL activation observed during the presentation of the aerial movie. These results suggest that the MTL is strongly engaged with encoding spatial information represented in multiple orientations, whereas particular perspectives (specifically, whether they are ground-level or aerial) in which the spatial information is depicted have little relevance to the degree of activation of MTL structures. Given the age-related atrophy and associated functional decline of the MTL discussed earlier, the neuroimaging findings pose the possibility that older adults find it equally difficult to learn environmental layouts from the ground-level and aerial-with-turns movies due to the need for processing varying orientations. Thus, in the experiment reported below, it was expected that older participants would form less accurate spatial memories following viewing the ground-level and aerial-with-turns movies than the aerial movie.

## 2. Material and methods

### 2.1 Participants

Eighteen younger and seventeen older adults were initially recruited for this study from the Brisbane community and students of Queensland University of Technology. Three of those older participants misunderstood instructions and generated non-interpretable data, and thus they were replaced with three new participants. As a result, the total of 38 participants took part in this study. They gave written informed consent prior to their participation in the study and received either partial course credit or monetary compensation.

Participants’ demographic characteristics are summarized in Table 1. The younger and older groups differed in the years of education they received, but there was no statistically significant difference between the groups in the ratios of male and female participants and in scores on the Saint Louis University mental status (SLUMS) examination [30], a standardized test for detecting dementia and mild cognitive impairment. The older group’s SLUMS scores were very similar to those of the younger group, suggesting that the older group represented the population of cognitively healthy older adults.

**Table 1.** Characteristics of Younger and Older Participant Groups

| | Younger ($n = 18$) | Older ($n = 17$) |
| --- | --- | --- |
| Age (years)* | 17–29<br>$M = 19.5, SD = 3.31$ | 60–92<br>$M = 70.5, SD = 8.48$ |
| Sex (male/female) | 5/13 | 6/11 |
| Education (years)* | 12–17<br>$M = 12.5, SD = 1.34$ | 10–20<br>$M = 14.76, SD = 2.61$ |
| SLUMS score (max = 30) | 24–30<br>$M = 27.67, SD = 1.81$ | 22–30<br>$M = 27.24, SD = 2.63$ |

*Note.* SLUMS = Saint Louis University mental status examination
\*The younger and older groups differed ( $p < .003$ ).

### 2.2 Design and procedure

The materials, design, and procedure of the current experiment were identical to those of the Yamamoto and DeGirolamo [7] study unless noted otherwise below. Importantly, this experiment used the aerial-with-turns movie in addition to the ground-level and aerial movies.

Three rectangular virtual environments (110 × 130 ft in virtual space) were presented on a desktop display using the PsychoPy program [31]. Each environment contained 10 large landmarks and 7 small objects in a unique configuration. These environments were originally constructed by Shelton and colleagues and used in their studies [8, 10, 11].

Each participant viewed all three environments. One of them was shown from the perspective of a six-foot-tall navigator on the ground (ground-level perspective), another from the perspective of an observer who was looking straight down the environment from the height of 70 feet in the north-up orientation (aerial perspective), and the other from the perspective of an aerial observer at the same height who turned 90° to the right at each corner of the environment (aerial-with-turns perspective). In each movie, only a section of the environment containing two to three landmarks was visible at any given time (Figure 1A–C). Across younger participants, the order of presentation of the three perspectives was counterbalanced, and each environment was presented in all three perspectives the equal number of times. The same design was used in the older group.

Participants were told that they would view movies of rectangular environments and learn locations of landmarks in each environment. They were informed in advance that the movies would start at the southwest corner and move along the four legs of an environment in clockwise order (Figure 1D). In the Yamamoto and DeGirolamo [7] study, each segment of a movie was preceded by a label that explicitly specified which leg participants were to experience. In the current experiment, these labels were removed and the four segments of the movie were presented as one continuous sequence that lasted approximately 40 s. During the first run-through of the movie, an experimenter pointed and named the 10 landmarks and clarified that their locations must be remembered later. Subsequently, participants viewed the movie six more times, during which they named each landmark as they saw it. After finishing the last run of the movie, participants were asked to mark the locations of the landmarks on a sheet of paper in which the perimeter of the environment was drawn to scale (16.8 × 19.9 cm). The southwest corner and the initial direction of travel were identified in this sheet. The names of the 10 landmarks were provided to participants. They were allowed to turn the sheet of paper in any direction while marking the landmark locations. They were given unlimited time to perform this task, and permitted to adjust marked landmark locations until they were satisfied with the map of the environment they created. They were not given any feedback on their performance. Once the map drawing task was completed for one environment, participants viewed the movie of the second environment and drew its map by following the same procedure. This was repeated for the third environment.

Accuracy of maps drawn by participants was quantified with Waterman and Gordon’s [32] distortion index (DI). In this approach, the layout of landmarks on a participant’s map was translated, rotated, and linearly scaled so that the difference between the drawn and true layouts of the landmarks was minimized. The optimal transformation of the drawn layout was uniquely determined by the least square method, and a DI was derived from the parameters of this transformation such that it represented the degree of distortion of the drawn layout independently of translation, rotation, and scaling factors. Theoretically, DIs can range from 0 (an accurately drawn layout) to 100 (a fully collapsed layout). Details of the definition and computational procedure of a DI are available elsewhere [32, 33].

In addition to the spatial learning tasks described above, all participants were given the SLUMS. One question in the SLUMS was modified so that it was suitable to Australian participants—the original version requires knowledge of American geography (i.e., Chicago is in Illinois), and thus this question was given using Melbourne and Victoria instead.

## 3. Results

DIs were analyzed by a mixed analysis of variance in which perspective (ground-level, aerial, and aerial-with-turns) was a within-participant factor and age (younger and older) was a between-participant factor. Greenhouse-Geisser correction was made for non-sphericity when appropriate.

Both the perspective hypothesis and the orientation hypothesis led to the prediction that the interaction between perspective and age would be significant. Thus, critical tests for distinguishing between the hypotheses were given by six planned contrasts in which the effects of each perspective were compared one-on-one by paired *t*-tests within each age group. These tests used a Bonferroni-corrected α of .008.

Figure 2 shows individual DIs of all participants as well as their means as a function of perspective and age. As clearly seen in the figure, DIs yielded from the ground-level and aerial-with-turns movies were similar to each other, and they were greater than those from the aerial movie on average. In line with this observation, the main effect of perspective was significant, *F*(2, 66) = 11.29, *p* < .001, η_G_^2^ = .14. Importantly, the effect of perspective was more pronounced in the older group than in the younger group, as shown by the significant interaction between perspective and age, *F*(2, 66) = 4.88, *p* = .02, η_G_^2^ = .07. In addition, the older group produced more distorted maps than the younger group regardless of perspectives, making the main effect of age significant, *F*(1, 33) = 23.90, *p* < .001, η_G_^2^ = .27.

**Figure 2.**
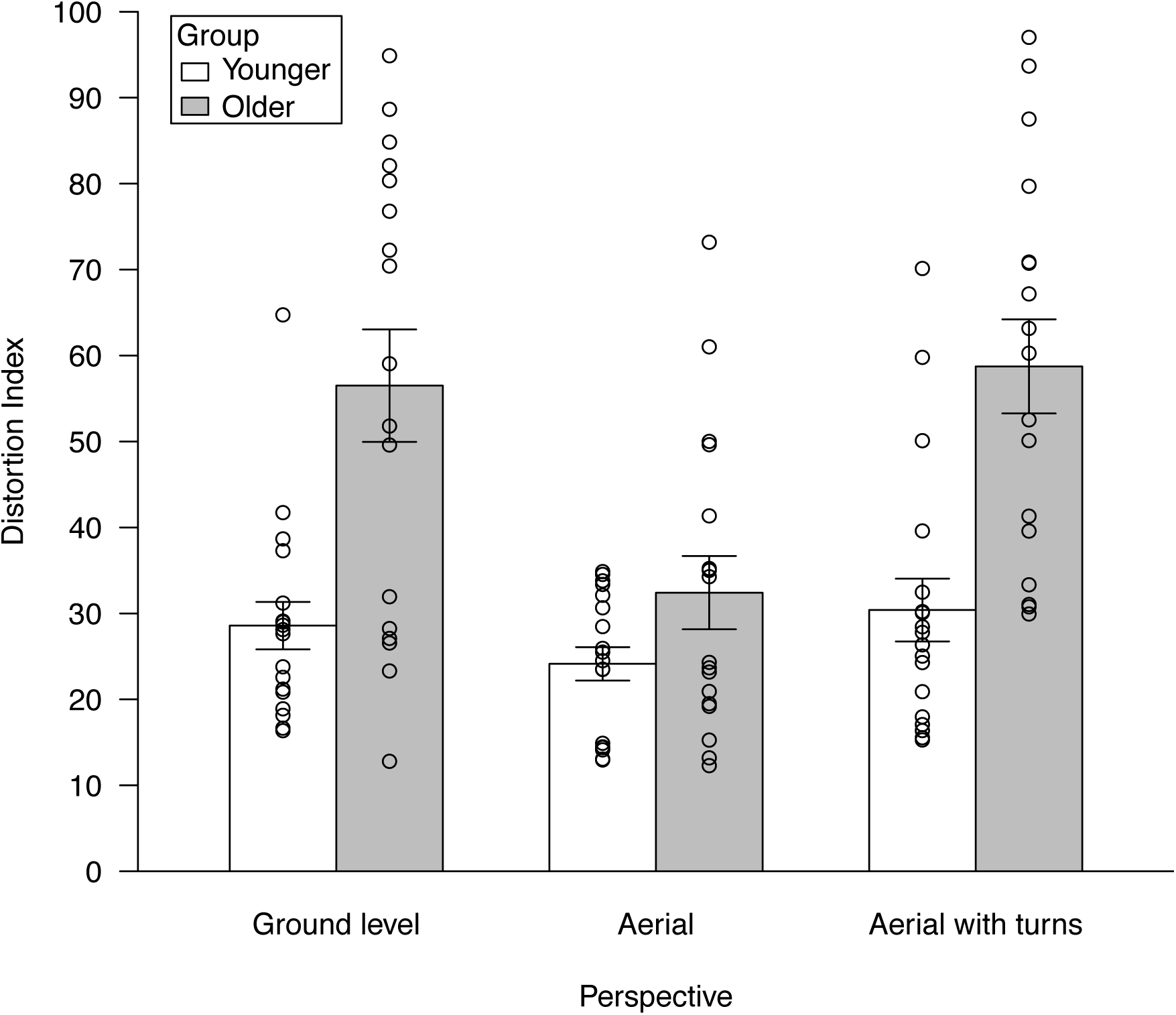
Distortion indices (DIs) of maps of learned environments produced by participants. Each open circle represents one participant and bars show mean DIs as a function of learning perspectives and age groups. Error bars indicate ±1 standard error of the mean.

One-on-one comparisons between perspectives within each age group corroborated the above observations. In the younger group, none of these comparisons reached significance—ground-level vs. aerial: *t*(17) = 1.33, *p* = .20, 95% CI [−2.62, 11.52]; ground-level vs. aerial-with-turns: *t*(17) = 0.47, *p* = .64, 95% CI [−6.28, 9.91]; and aerial vs. aerial-with-turns: *t*(17) = 1.61, *p* = .13, 95% CI [−1.94, 14.46]. On the other hand, in the older group, while mean DIs from the ground-level and aerial-with-turns movies did not differ from each other, *t*(16) = 0.28, *p* = .79, 95% CI [−14.98, 19.46], they were significantly greater than those from the aerial movie, *t*(16) = 4.66, *p* < .001, 95% CI [13.12, 35.05] (ground-level vs. aerial) and *t*(16) = 4.62, *p* < .001, 95% CI [14.23, 38.41] (aerial vs. aerial-with-turns).

In sum, accuracy of younger participants’ memories for landmark layouts was not affected by perspectives from which environments were viewed, whereas an equal degree of distortion was introduced in older participants’ memories for the layouts when they were learned from ground-level and aerial-with-turns movies—that is, when the layouts were encoded in the presence of orientation changes regardless of the perspectives.

## 4. Discussion

Results of the current experiment were very clear about the perspective and orientation hypotheses: The former was rejected and the latter was supported. Older participants built less accurate memories for large-scale environments when they viewed different portions of the environments from different orientations. Importantly, unlike in the previous study [7], the presence of varying orientations during learning was not unique to a particular perspective. Instead, in this study, it was present both in the ground-level movie and in the aerial-with-turns movie. Furthermore, relative to older participants’ memories formed through the fixed-orientation aerial movie, their memories encoded via the ground-level and aerial-with-turns movies showed the same degree of additional distortion. Together, these findings indicate that ground-level vs. aerial perspectives per se had minimal influence on older participants’ performance in spatial learning. By contrast, the difficulty of spatial learning was disproportionately increased in the older group when a memory for the entire environment had to be constructed from spatial information that was initially represented in divergent orientations. Critically, these patterns were virtually absent in younger participants (i.e., they performed similarly in all conditions), suggesting that the salient effect of orientation change during environmental learning on subsequent spatial memory is a consequence of cognitive and neurological aging processes.

As discussed in the introduction, empirical support for the orientation hypothesis was predicted in this study on the basis of previous neuroimaging findings that the MTL is particularly engaged with environmental learning that entails processing of orientation changes [10, 11]. These findings were obtained largely from younger participants (e.g., the mean age of participants was 23.1 years in [10]). The current results extend the previous findings by showing that increased distortion of spatial memory follows selectively from encoding conditions involving multiple orientations, when MTL functionality is presumably deteriorated due to old age [12–17]. Although this functional decline is only assumed in the present study, Borghesani et al. [9] demonstrated that older adults who were carriers of the apolipoprotein ε4 allele, a risk factor for Alzheimer’s disease that alters MTL activation even before the onset of disease symptoms [34–36], exhibited reduced blood-oxygen-level-dependent signals in the MTL while learning a large-scale environment via the ground-level movie (as compared to age-matched participants who were not carriers of the allele). In other words, older adults who had genetic evidence for declining MTL functions actually showed an altered level of MTL activation in response to spatial learning with multiple orientations, suggesting that functional alteration of the MTL most likely underlies the observed pattern of older participants’ performance in the present study.

Considering that the MTL carries out various processes of spatial learning and memory [37, 38], a question remains as to which of those processes were most critically affected by aging and thus primarily accounted for the selective impairment in encoding an environmental layout from multiple orientations. To address this question, it is useful to examine which non-MTL areas of the brain show activation patterns that co-vary with those of the MTL as a function of the three types of movies. In the Shelton and Pippitt [11] study, the right inferior parietal cortex (Brodmann area [BA] 40) and the right posterior cingulate cortex (BA 31) exhibited the same level of activation in response to the ground-level and aerial-with-turns movies, which was greater than the level of activation in response to the aerial movie. Activation in the inferior parietal cortex indicates that the ground-level and aerial-with-turns movies indeed required participants to mentally rotate the views of an environment, as there is accumulating evidence that this area is involved in mental rotation [39–42]. Previous studies also showed that posterior cingulate neurons are responsive to navigators’ headings in a large-scale environment (those in BA 31 [43]) and considered to take part in translating spatial information between egocentric and allocentric frames of reference (those in BA 29 and 30; i.e., the retrosplenial cortex [38, 44–46]). Activation of these neurons suggests that participants initially encoded the views of the environment in a heading-dependent manner, and then converted them into a common frame of reference. Given the dense projection from the posterior cingulate region to the MTL [44] and the MTL’s well-defined role in building mental representations of surrounding environments (i.e., the so-called cognitive maps [38, 47]), it is inferred that the main function of the MTL in the current paradigm was to receive input from the posterior cingulate region and construct a unified representation of the entire environment by completing the conversion of the views. This integrative function of the MTL may be susceptible to age-related decline, leading to formation of less integrated and thus more distorted memories of large-scale environments. In addition, this decline may be exacerbated by deteriorated input into the MTL (i.e., inadequately performed conversion of spatial information between reference frames) due to metabolic alteration of the posterior cingulate region in the aged brain [48, 49]. To refine and support these inferences, a neuroimaging study should be conducted in the future for directly contrasting activation in the MTL and posterior cingulate region while older adults learn large-scale environments with fixed and dynamic orientations.

Shelton and Pippitt [11] also showed that several areas of the brain were activated by the aerial and aerial-with-turns movies in the same manner. They included the superior parietal cortex (BA 7), fusiform gyrus (BA 19 and 37), left medial frontal cortex (BA 6), and right middle occipital gyrus (BA 18). It is likely that these areas responded to the aerial perspective itself regardless of whether movies contained changing orientations. In fact, past research suggests that activation in many of these areas was primarily driven by pictorial properties of the movies, not by information about environmental layouts that the movies provided. For example, the superior parietal cortex is involved in representing space around one’s body, and a critical factor that determines the locus of neural processing in this brain area is the distance between the represented space and the body [50–52]. Given that the two types of aerial movies depicted environments from the same height, it is reasonable that they elicited comparable activation patterns in this area. The fusiform and middle occipital gyri are visual areas of the brain that mainly participate in object and face processing [53–55]. It was probable that these areas were activated by the appearance of landmarks, which was similar between the aerial and aerial-with-turns movies. Older participants in the present study performed well in forming memories for the environments via the aerial movie, suggesting that functions of these brain areas relevant to spatial learning (e.g., visually recognizing individual landmarks) are preserved relatively well in healthy aging process.

Finally, it is worth mentioning that the present results addressed another limitation of the previous study [7] caused by the use of the map drawing task. Although this task was effective for assessing memory for a spatial layout as a whole, it could have given spatial learning via the aerial movie an advantage because the map drawing task required participants to produce aerial views of environmental layouts. That is, only after learning an environment via the ground-level movie, participants needed to mentally convert their spatial memory from ground-level to aerial perspectives. Thus, reduced accuracy of participants’ maps in the ground-level condition could have been caused by this post-encoding transformation of perspectives, not by orientations or perspectives that participants had during memory encoding. This possibility is now ruled out because spatial learning via the aerial-with-turns movie did not entail the post-encoding perspective conversion, but it still resulted in participants’ maps that contained the same degree of distortion as those yielded from the ground-level movie. It can be concluded that effects of perspectives were negligible in the current paradigm, regardless of whether they came from processing of different views during learning or transformation of memory representations from one perspective to another.

## 5. Conclusions

Building upon the previous finding that age-related decline of environmental learning ability is more evident in first-person navigation than in map reading [7], the present study investigated what aspects of first-person navigation make this type of spatial learning particularly challenging to older adults. Specifically, it was hypothesized that either the ground-level perspective that navigators take during navigation or varying orientations from which navigators view an environment would increase the difficulty of encoding the environment through navigation. To distinguish between these two possibilities, this study added a new learning condition in which participants had aerial views of the environment that rotated 90° at each corner. In this condition, healthy older participants were as impaired as in the ground-level condition, while they formed significantly more accurate memory for the environment from fixed-orientation aerial images. These results suggest that the source of older adults’ spatial learning impairment is not a particular perspective (i.e., ground-level or aerial) but the need for processing multiple orientations of the environment during memory formation. In the brain, it is likely that this impairment is attributed to the MTL, and possibly to the posterior cingulate region as well. Future research should refine the understanding of functional alteration of these areas in the aged brain so that older adults’ difficulty with spatial learning and navigation can be more precisely characterized.

## Acknowledgements

This research was supported by Early Career Academic Recruitment and Development Grant from Queensland University of Technology and Faculty Research and Development Award from Cleveland State University. The funders had no role in study design, data collection and analysis, decision to publish, or preparation of the manuscript. The authors thank Amy Shelton for providing stimuli used in this research.

## References

[1] P.C. Burns, Navigation and the mobility of older drivers, J. Gerontol. B. Psychol. Sci. Soc. Sci. 54B (1999) S49–S55. https://doi.org/10.1093/geronb/54B.1.S49.

[2] S.D. Moffat, Aging and spatial navigation: What do we know and where do we go?, Neuropsychol. Rev. 19 (2009) 478–489. https://doi.org/10.1007/s11065-009-9120-3.

[3] M.-C. Pai, W.J. Jacobs, Topographical disorientation in community-residing patients with Alzheimer’s disease, Int. J. Geriatr. Psychiatry. 19 (2004) 250–255. https://doi.org/10.1002/gps.1081.

[4] A.E. Sanders, R. Holtzer, R.B. Lipton, C. Hall, J. Verghese, Egocentric and exocentric navigation skills in older adults, J. Gerontol. A. Biol. Sci. Med. Sci. 63 (2008) 1356–1363. https://doi.org/10.1093/gerona/63.12.1356.

[5] M.A. Harris, J.M. Wiener, T. Wolbers, 2012. Aging specifically impairs switching to an allocentric navigational strategy, Front. Aging Neurosci. 4, 29. https://doi.org/10.3389/fnagi.2012.00029.

[6] V. Muffato, C. Meneghetti, R. De Beni, Not all is lost in older adults’ route learning: The role of visuo-spatial abilities and type of task, J. Environ. Psychol. 47 (2016) 230–241. https://doi.org/10.1016/j.jenvp.2016.07.003.

[7] N. Yamamoto, G.J. DeGirolamo, 2012. Differential effects of aging on spatial learning through exploratory navigation and map reading, Front. Aging Neurosci. 4, 14. https://doi.org/10.3389/fnagi.2012.00014.

[8] A.L. Shelton, T.P. McNamara, Orientation and perspective dependence in route and survey learning, J. Exp. Psychol. Learn. Mem. Cogn. 30 (2004) 158–170. https://doi.org/10.1037/0278-7393.30.1.158.

[9] P.R. Borghesani, L.C. Johnson, A.L. Shelton, E.R. Peskind, E.H. Aylward, G.D. Schellenberg, M.M. Cherrier, Altered medial temporal lobe responses during visuospatial encoding in healthy APOE*4 carriers, Neurobiol. Aging. 29 (2008) 981–991. https://doi.org/10.1016/j.neurobiolaging.2007.01.012.

[10] A.L. Shelton, J.D.E. Gabrieli, Neural correlates of encoding space from route and survey perspectives, J. Neurosci. 22 (2002) 2711–2717. https://doi.org/10.1523/JNEUROSCI.22-07-02711.2002.

[11] A.L. Shelton, H.A. Pippitt, Fixed versus dynamic orientations in environmental learning from ground-level and aerial perspectives, Psychol. Res. 71 (2007) 333–346. https://doi.org/10.1007/s00426-006-0088-9.

[12] J. Golomb, M.J. de Leon, A. Kluger, A.E. George, C. Tarshish, S.H. Ferris, Hippocampal atrophy in normal aging: an association with recent memory impairment, Arch. Neurol. 50 (1993) 967–973. https://doi.org/10.1001/archneur.1993.00540090066012.

[13] T.L. Jernigan, S.L. Archibald, C. Fennema-Notestine, A.C. Gamst, J.C. Stout, J. Bonner, J.R. Hesselink, Effects of age on tissues and regions of the cerebrum and cerebellum, Neurobiol. Aging. 22 (2001) 581–594. https://doi.org/10.1016/S0197-4580(01)00217-2.

[14] S.D. Moffat, W. Elkins, S.M. Resnick, Age differences in the neural systems supporting human allocentric spatial navigation, Neurobiol. Aging. 27 (2006) 965–972. https://doi.org/10.1016/j.neurobiolaging.2005.05.011.

[15] Z. Nedelska, R. Andel, J. Laczó, K. Vlcek, D. Horinek, J. Lisy, K. Sheardova, J. Bureš, J. Hort, Spatial navigation impairment is proportional to right hippocampal volume, Proc. Natl. Acad. Sci. 109 (2012) 2590–2594. https://doi.org/10.1073/pnas.1121588109.

[16] R.I. Scahill, C. Frost, R. Jenkins, J.L. Whitwell, M.N. Rossor, N.C. Fox, A longitudinal study of brain volume changes in normal aging using serial registered magnetic resonance imaging, Arch Neurol. 60 (2003) 989–994. https://doi.org/10.1001/archneur.60.7.989.

[17] I. Driscoll, D.A. Hamilton, H. Petropoulos, R.A. Yeo, W.M. Brooks, R.N. Baumgartner, R.J. Sutherland, The aging hippocampus: cognitive, biochemical and structural Findings, Cereb. Cortex. 13 (2003) 1344–1351. https://doi.org/10.1093/cercor/bhg081.

[18] J.M. Loomis, J.A. Da Silva, N. Fujita, S.S. Fukusima, Visual space perception and visually directed action, J. Exp. Psychol. Hum. Percept. Perform. 18 (1992) 906–921. https://doi.org/10.1037/0096-1523.18.4.906.

[19] R.C. Toye, The effect of viewing position on the perceived layout of space, Percept. Psychophys. 40 (1986) 85–92. https://doi.org/10.3758/BF03208187.

[20] M. Wagner, The metric of visual space, Percept. Psychophys. 38 (1985) 483–495. https://doi.org/10.3758/BF03207058.

[21] Z. Bian, G.J. Andersen, Aging and the perception of egocentric distance, Psychol. Aging. 28 (2013) 813–825. https://doi.org/10.1037/a0030991.

[22] J.F. Norman, O.C. Adkins, H.F. Norman, A.G. Cox, C.E. Rogers, Aging and the visual perception of exocentric distance, Vision Res. 109 (2015) 52–58. https://doi.org/10.1016/j.visres.2015.02.007.

[23] J.F. Norman, O.C. Adkins, L.E. Pedersen, C.M. Reyes, R.A. Wulff, A. Tungate, The visual perception of exocentric distance in outdoor settings, Vision Res. 117 (2015) 100–104. https://doi.org/10.1016/j.visres.2015.10.003.

[24] J.E. Birren, L.M. Fisher, Aging and speed of behavior: possible consequences for psychological functioning, Annu. Rev. Psychol. 46 (1995) 329–353. https://doi.org/10.1146/annurev.ps.46.020195.001553.

[25] T.A. Salthouse, The processing-speed theory of adult age differences in cognition, Psychol. Rev. 103 (1996) 403–428. https://doi.org/10.1037/0033-295X.103.3.403.

[26] S.A. Gaylord, G.R. Marsh, Age differences in the speed of a spatial cognitive process, J. Gerontol. 30 (1975) 674–678. https://doi.org/10.1093/geronj/30.6.674.

[27] J.F. Herman, P.R. Bruce, Adults’ mental rotation of spatial information: effects of age, sex and cerebral laterality, Exp. Aging Res. 9 (1983) 83–85. https://doi.org/10.1080/03610738308258430.

[28] C. Hertzog, B. Rypma, Age differences in components of mental-rotation task performance, Bull. Psychon. Soc. 29 (1991) 209–212. https://doi.org/10.3758/BF03335237.

[29] J.T. Puglisi, R.W. Morrell, Age-related slowing in mental rotation of three-dimensional objects, Exp. Aging Res. 12 (1986) 217–220. https://doi.org/10.1080/03610738608258571.

[30] S.H. Tariq, N. Tumosa, J.T. Chibnall, M.H. Perry, J.E. Morley, Comparison of the Saint Louis University Mental Status examination and the mini-mental state examination for detecting dementia and mild neurocognitive disorder—a pilot study, Am. J. Geriatr. Psychiatry. 14 (2006) 900–910. https://doi.org/10.1097/01.JGP.0000221510.33817.86.

[31] J.W. Peirce, PsychoPy—psychophysics software in Python, J. Neurosci. Methods. 162 (2007) 8–13. https://doi.org/10.1016/j.jneumeth.2006.11.017.

[32] S. Waterman, D. Gordon, A quantitative-comparative approach to analysis of distortion in mental maps., Prof. Geogr. 36 (1984) 326–337. https://doi.org/10.1111/j.0033-0124.1984.00326.x.

[33] W.R. Tobler, Bidimensional regression, Geogr. Anal. 26 (1994) 187–212. https://doi.org/10.1111/j.1538-4632.1994.tb00320.x.

[34] M.A. Trivedi, T.W. Schmitz, M.L. Ries, B.M. Torgerson, M.A. Sager, B.P. Hermann, S. Asthana, S.C. Johnson, 2006. Reduced hippocampal activation during episodic encoding in middle-aged individuals at genetic risk of Alzheimer’s Disease: a cross-sectional study, BMC Med. 4, 1. https://doi.org/10.1186/1741-7015-4-1.

[35] N.A. Dennis, J.N. Browndyke, J. Stokes, A. Need, J.R. Burke, K.A. Welsh-Bohmer, R. Cabeza, Temporal lobe functional activity and connectivity in young adult *APOE* ε4 carriers, Alzheimers Dement. J. Alzheimers Assoc. 6 (2010) 303–311. https://doi.org/10.1016/j.jalz.2009.07.003.

[36] N. Filippini, B.J. MacIntosh, M.G. Hough, G.M. Goodwin, G.B. Frisoni, S.M. Smith, P.M. Matthews, C.F. Beckmann, C.E. Mackay, Distinct patterns of brain activity in young carriers of the *APOE*-ε4 allele, Proc. Natl. Acad. Sci. 106 (2009) 7209–7214. https://doi.org/10.1073/pnas.0811879106.

[37] N. Burgess, E.A. Maguire, J. O’Keefe, The human hippocampus and spatial and episodic memory, Neuron. 35 (2002) 625–641. https://doi.org/10.1016/S0896-6273(02)00830-9.

[38] R.A. Epstein, E.Z. Patai, J.B. Julian, H.J. Spiers, The cognitive map in humans: spatial navigation and beyond, Nat. Neurosci. 20 (2017) 1504–1513. https://doi.org/10.1038/nn.4656.

[39] P.A. Carpenter, M.A. Just, T.A. Keller, W. Eddy, K. Thulborn, Graded functional activation in the visuospatial system with the amount of task demand, J. Cogn. Neurosci. 11 (1999) 9–24. https://doi.org/10.1162/089892999563210.

[40] M.S. Cohen, S.M. Kosslyn, H.C. Breiter, G.J. DiGirolamo, W.L. Thompson, A.K. Anderson, S.Y. Brookheimer, B.R. Rosen, J.W. Belliveau, Changes in cortical activity during mental rotation: a mapping study using functional MRI, Brain 119 (1996) 89–100. https://doi.org/10.1093/brain/119.1.89.

[41] A.L. Shelton, H.A. Pippitt, Motion in the mind’s eye: comparing mental and visual rotation, Cogn. Affect. Behav. Neurosci. 6 (2006) 323–332. https://doi.org/10.3758/CABN.6.4.323.

[42] J. Vanrie, E. Béatse, J. Wagemans, S. Sunaert, P. Van Hecke, Mental rotation versus invariant features in object perception from different viewpoints: an fMRI study, Neuropsychologia. 40 (2002) 917–930. https://doi.org/10.1016/S0028-3932(01)00161-0.

[43] O. Baumann, J.B. Mattingley, Medial parietal cortex encodes perceived heading direction in humans, J. Neurosci. 30 (2010) 12897–12901. https://doi.org/10.1523/JNEUROSCI.3077-10.2010.

[44] S.D. Vann, J.P. Aggleton, E.A. Maguire, What does the retrosplenial cortex do?, Nat. Rev. Neurosci. 10 (2009) 792–802. https://doi.org/10.1038/nrn2733.

[45] S.A. Marchette, L.K. Vass, J. Ryan, R.A. Epstein, Anchoring the neural compass: coding of local spatial reference frames in human medial parietal lobe, Nat. Neurosci. 17 (2014) 1598–1606. https://doi.org/10.1038/nn.3834.

[46] P. Byrne, S. Becker, N. Burgess, Remembering the past and imagining the future: a neural model of spatial memory and imagery., Psychol. Rev. 114 (2007) 340–375. https://doi.org/10.1037/0033-295X.114.2.340.

[47] C.M. Bird, N. Burgess, The hippocampus and memory: insights from spatial processing, Nat Rev Neurosci. 9 (2008) 182–194. https://doi.org/10.1038/nrn2335.

[48] N. Villain, B. Desgranges, F. Viader, V. de la Sayette, F. Mézenge, B. Landeau, J.-C. Baron, F. Eustache, G. Chételat, Relationships between hippocampal atrophy, white matter disruption, and gray matter hypometabolism in Alzheimer’s disease, J. Neurosci. 28 (2008) 6174–6181. https://doi.org/10.1523/JNEUROSCI.1392-08.2008.

[49] K. Mevel, B. Desgranges, J.-C. Baron, B. Landeau, V. De la Sayette, F. Viader, F. Eustache, G. Chételat, Detecting hippocampal hypometabolism in Mild Cognitive Impairment using automatic voxel-based approaches, NeuroImage. 37 (2007) 18–25. https://doi.org/10.1016/j.neuroimage.2007.04.048.

[50] M. Corbetta, G.L. Shulman, Control of goal-directed and stimulus-driven attention in the brain, Nat. Rev. Neurosci. 3 (2002) 201–215. https://doi.org/10.1038/nrn755.

[51] J. Cléry, O. Guipponi, C. Wardak, S. Ben Hamed, Neuronal bases of peripersonal and extrapersonal spaces, their plasticity and their dynamics: knowns and unknowns, Neuropsychologia. 70 (2015) 313–326. https://doi.org/10.1016/j.neuropsychologia.2014.10.022.

[52] P.W. Halligan, J.C. Marshall, Left neglect for near but not far space in man, Nature. 350 (1991) 498–500. https://doi.org/10.1038/350498a0.

[53] L.G. Ungerleider, A.H. Bell, Uncovering the visual “alphabet”: advances in our understanding of object perception, Vision Res. 51 (2011) 782–799. https://doi.org/10.1016/j.visres.2010.10.002.

[54] J.V. Haxby, M.I. Gobbini, M.L. Furey, A. Ishai, J.L. Schouten, P. Pietrini, Distributed and overlapping representations of faces and objects in ventral temporal cortex, Science. 293 (2001) 2425–2430. https://doi.org/10.1126/science.1063736.

[55] J. Sergent, S. Ohta, B. MacDonald, Functional neuroanatomy of face and object processing: a positron emission tomography study, Brain. 115 (1992) 15–36. https://doi.org/10.1093/brain/115.1.15.

